# E-cadherin biointerfaces reprogram collective cell migration and cell cycling by forcing homeostatic conditions

**DOI:** 10.1101/2023.07.25.550505

**Authors:** Kevin Suh, Youn Kyoung Cho, Isaac B. Breinyn, Daniel J. Cohen

## Abstract

Cells attach to the world around them in two ways—cell:extracellular-matrix adhesion and cell:cell adhesion—and conventional biomaterials are made to resemble the matrix to encourage integrin-based cell adhesion. However, interest is growing for cell-mimetic interfaces that mimic cell-cell interactions using cadherin proteins, as this offers a new way to program cell behavior and design synthetic implants and objects that can integrate directly into living tissues. Here, we explore how these cadherin-based materials affect collective cell behaviors, focusing specifically on collective migration and cell cycle regulation in cm-scale epithelia. We built culture substrates where half of the culture area was functionalized with matrix proteins and the contiguous half was functionalized with E-cadherin proteins, and we grew large epithelia across this ‘Janus’ interface. Parts of the tissues in contact with the matrix side of the Janus interface exhibited normal collective dynamics, but an abrupt shift in behaviors happened immediately across the Janus boundary onto the E-cadherin side, where cells formed hybrid E-cadherin junctions with the substrate, migration effectively froze in place, and cell-cycling significantly decreased. E-cadherin materials suppressed long-range mechanical correlations in the tissue and mechanical information reflected off the substrate interface. These effects could not be explained by conventional density, shape index, or contact inhibition explanations. E-cadherin surfaces nearly doubled the length of the G0/G1 phase of the cell cycle, which we ultimately connected to the exclusion of matrix focal adhesions induced by the E-cadherin culture surface.

## Introduction

There are two ways that cells in multicellular organisms can attach to the world around themselves–cell-substrate adhesion and cell-cell adhesion (1, 2). Manipulating where and how cells can attach to their surroundings is a powerful strategy for understanding cell biology, characterizing cellular biophysics, and designing biomaterial interfaces. Broadly, cell-substrate adhesion relies on integrin protein-mediated attachments through focal adhesions—’foot’ complexes that allow cells to bind to the non-living extracellular matrix (ECM) around them such as collagen and fibronectin (3). By coupling the contractile cytoskeleton to these substrate adhesions, cells can then pull on the world around themselves to change shape or move and to sense the mechanical properties of their environment (4, 5). Cell-cell adhesion, by contrast, allows cells to attach to, and pull on, each other through a completely different set of cell-cell adhesion proteins that include cadherins, CAMs, and nectins (2, 6, 7). Cell-cell adhesions serve multiple mechanical functions: tissue integrity (e.g. E-cadherin and desmosomes); barrier function (e.g. tight junctions); mechanotransduction of cell-cell forces (e.g. cadherins); and general cell-cell recognition, making cell-cell adhesions akin to ‘handshakes’ (6). While cell-substrate and cell-cell adhesions are both critical, the majority of work on cell-material interfaces has targeted cell-substrate, ‘foot-like’ interactions, from the earliest Egyptian microfiber scaffolds (c. 5000 BCE) to modern hydrogel and decellularized-ECM biomaterials. How might materials that deliberately target cell-cell adhesion and cellular ‘handshakes’ instead of cell-substrate ‘foot’ interactions change cell-material interactions? There is significant interest in this question given the broad importance of cell-cell adhesions to biological and biophysical processes, along with a growing need for new paradigms in cell-material attachment for biotechnology and biomedical applications. Here, we specifically focus on how such cell-mimetic interfaces can regulate and reprogram large-scale, multicellular collective cell behavior in patterned epithelial tissue layers.

How can we make cell-mimetic interfaces that specifically target cell-cell interactions? In the same way that functionalizing any arbitrary synthetic material with ECM proteins (e.g. collagen, fibronectin, laminin, etc.) or ECM-mimics (e.g. RGD peptide) causes cells to attach to them via integrin/foot-processes, functionalizing a material with cell-cell adhesion proteins (e.g. purified extra-cellular domains of cadherins (8–10)) or peptide mimics (e.g. HAV) (11) can trigger cellular processes and pathways normally only active when a cell meets another cell. Further, just as the community has thoroughly characterized substrate-like biomaterials for how ECM stiffness (e.g. soft vs. hard (12–14)), viscoelasticity (15), and composition (e.g. collagen, fibronectin, etc.) regulate cell behaviors, the same approach is gradually being applied to cell-mimetic interfaces with applications spanning stem cell maintenance (16), reductionist assays for cell junction and signaling biology (17–19), and early approaches for improved stability of cell-material interfaces in 2D (20) and 3D (8) for eventual translational applications (21, 22). Interestingly, a key area of study for ECM substrates—how they regulate dynamic collective cell behaviors in large groups of cells such as migration and proliferation(23–25)—has been little explored with cell-mimetic surfaces, where the focus has more often been on single cell biology or static endpoint assays with fixed tissues (11, 18, 20), leaving many open questions. Collective and coordinated cell behaviors are absolutely critical to multicellular, and cadherins themselves play a key role in many cases such as transmitting tension information, maintaining tissue, and enabling large-scale crowd dynamics during healing and development (26, 27), while misregulation of cadherin adhesions is implicated in a number of cancers (28, 29). Given this key role for cadherins in collective cell behaviors, we hypothesized that an E-cadherin-based culture substrate would dramatically alter collective dynamics compared to traditional culture substrates.

To best capture differences in collective cell dynamics on cell-mimetic vs. substrate-mimetic interfaces, we created a hybrid culture surface where one-half of the surface was functionalized to resemble ECM and target cellular ‘feet’ (integrins), and the other half was functionalized with purified, extracellular E-cadherin (Ecad:Fc) to target cellular ‘handshakes’. Such materials with two functional ‘faces’ are called Janus materials after the two-faced Greek god (30), and we cultured large, continuous epithelial layers across the boundaries of our Janus surfaces and then performed timelapse microscopy to investigate how these different surfaces affected collective cell behaviors. We used Madin-Darby canine kidney epithelium (MDCK) as it is the gold standard for collective cell dynamics studies (31–33) and exhibits extremely well-characterized native E-cadherin junctions as well as strong attachments to functionalized Ecad:Fc substrates (8, 10). Continuous epithelia grown on Janus surfaces sharply transitioned from integrin-adhesions on the ECM side of the surface to basal cadherin attachments on the cell-mimetic side, with unique cytoskeletal architectures characterizing each interface, among other differences. Surprisingly, rather than upregulating E-cadherin we found that the tissue appears to maintain cadherin levels at a fixed amount, instead redistributing available E-cadherin between cell-cell junctions and the cell-mimetic surface. Such hybrid junctions to the material imposed dramatic consequences on collective migration, resulting in nearly 10X slower cell migration speed with significantly reduced cell-cell coordination. In fact, our data indicated that mechanical information could no longer propagate through cells attached to the cell-mimetic surface. However, actual morphological differences between cells were relatively subtle for the first few cell cycles within compared tissues. Cell proliferation itself was also markedly slowed, and we traced the origins of this to a nearly 2X increase specifically in the G0/G1 phase of the cell cycle of cells cultured on Ecad:Fc. These effects did not appear to be induced by jamming transitions anticipated in previous studies (34, 35), but instead appeared directly related to the lack of integrin-based adhesions on the cell-mimetic parts of the surface, hence even tissues with relatively similar morphologies can exhibit dramatically different collective dynamics based on how they attach to the world around themselves.

## Results

### Preparation and validation of ECM/Cadherin Janus substrates

We created the Janus substrates using a sequential stencil strategy where we used a silicone stencil to mask part of a glass culture surface and functionalized the exposed region of the substrate with collagen or cadherin. Then we removed the stencil and functionalized the fresh surface with the alternate protein—this ‘trapdoor’ process is shown in Figure 1A, B. Functionalizing the glass with an aldehyde silane first created an amine-reactive surface for protein crosslinking. Collagen-IV (Col-IV; 50 μg/mL) was directly coupled to this surface for the ECM side of the surface, while we use Protein-AG (PAG; 250 μg/mL) on the cell-mimetic surface to help properly orient Fc:E-cadherin (Ecad:Fc; 25 μg/mL), as in Fig. 1B. Here, we used purified human Fc:E-cadherin, a fusion of the Fc domain and the extracellular domains EC1-5 of E-cadherin (36), which is known to work best when properly oriented relative to the substrate (37). We used ELISA to determine the optimum Ecad:Fc concentration (Fig. S1). Fig. S2 shows a representative image of a Janus substrate with labeled ECM (green) and Ecad:Fc (magenta; antibody staining). Note that we did not fluorescently label the Ecad:Fc for any other data, so all fluorescence E-cadherin data in the paper comes from the cells themselves. See Methods for a full accounting, as well as prior related protocols. (8).

**Fig. 1.**
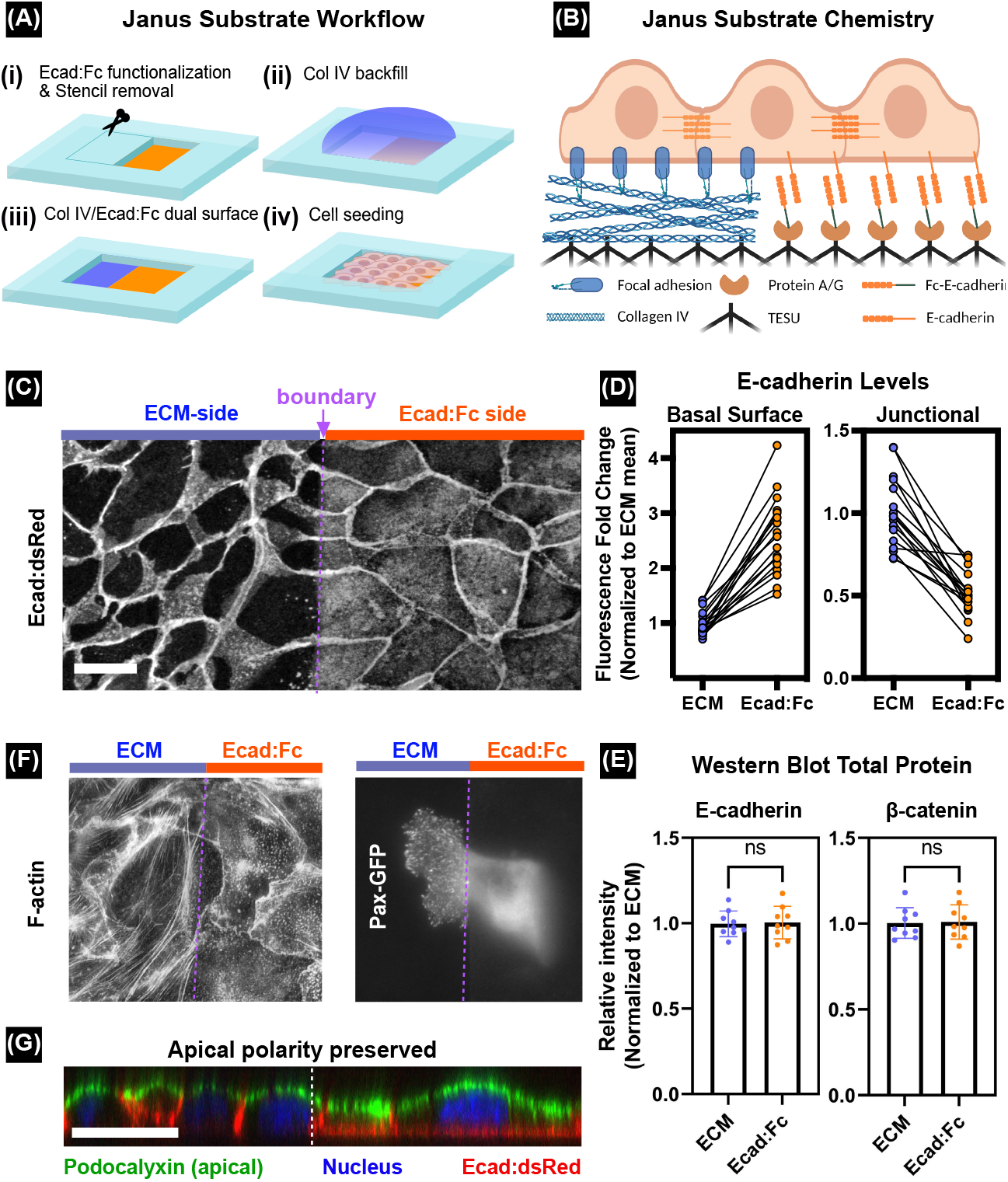
Functionalization and validation of ECM/Ecad:Fc Janus substrate. (A) Janus substrate functionalization strategy. (i) Ecad:Fc functionalization in the exposed region, while masking part of surface. Removal of the silicone stencil that covered the non-functionalized half to (ii) backfill with the Col IV solution. (iii) Preparation of contiguous Col IV/Ecad:Fc substrate. (iv) Monolayer of MDCK tissue seeded within the stencil. (B) Summary of Janus substrate surface chemistry. Focal adhesion formation and basal E-cadherin recruitment at the tissue-ECM interface and tissue-Ecad:Fc interface, respectively. Figure was generated with BioRender. Ecad:dsRed expressing MDCK monolayer on Janus boundary; ECM (left, blue), Ecad:Fc (right, orange), and dashed boundary (purple). (D) Quantification of the normalized average basal (left) and junctional (right) Ecad:dsRed signal across the boundary (n=17 across 3 experiments). (E) Quantification of Western blot analysis to detect the E-cadherin (left) and *β*-catenin (right) expression level (n=9 across 3 experiments). (F) *left:* Fluorescent labeling of MDCK tissue on Janus substrate with phalloidin to visualize F-actin cytoskeleton. *right:* Paxilin-GFP MDCK on Janus boundary. (G) Confocal XZ section of Ecad:dsRed MDCK on the Janus boundary, immunostained for an apical marker (podocalyxin) and nucleus (Hoechst). P-values are calculated using the unpaired, non-parametric Mann–Whitney test. ns: non-significant, Scale bar: 50 μm

A functional Janus surface should recruit cellular E-cadherin junctions to the basal plane of those cells in contact with the Ecad:Fc side. We validated and visualized this using Ecad:dsRed MDCK cells, noting that cells straddling the ECM/Fc-Ecad boundary actually showed split identities, with half of the cell’s basal surface recruiting E-cadherin (Fig. 1C) while the other half did not. We quantified the average intensity of these ‘hybrid junctions’ relative to the background basal signal on the ECM side of the Janus substrate and observed ∼2.5X increase in the basal E-cadherin signal on the Ecad:Fc substrate (Fig. 1D, left). Movies of this interface demonstrate both the sharpness and stability of the recruited E-cadherin (Movie S1). In contrast to basal E-cadherin increasing on the cell-mimetic substrate, junctional E-cadherin on average went down relative to that found in cells on standard ECM substrates (∼0.5X), suggesting a possible homeostatic balancing of the internal E-cadherin pool (Fig. 1D, right). We investigated this further using Western blotting (Fig. 1E, left) where we compared total E-cadherin levels in tissues grown purely on Ecad:Fc surfaces to similar tissues grown purely on ECM substrates and observed no statistical difference (see Fig. S3 for gel data). To further validate this, we also blotted for total *β*-catenin levels (key for the cadherin-catenin complex), again showing no apparent up- or downregulation (Fig. 1E, right). These data support the hypothesis that recruitment of cellular E-cadherin to the basal surface of cells on Ecad:Fc substrates may be balanced by depleted junctional E-cadherin.

E-cadherin plays an important role in cytoskeletal organization and cytoskeletal tension through the classical cadherin-catenin complex. Also, a previous study reported that the single cells cultured on cadherin substrates exhibit altered cytoskeletons (18), so we investigated the effects on junctioned tissue using an F-actin stain (Phalloidin, SI Appendix). Again, we observed a stark difference in cytoskeletal structures across the interface, with cells on ECM surfaces exhibiting the classic ‘stress fibers’ and junctional F-actin, while cells on the E-cadherin substrate exhibited many small F-actin puncta and junctional F-actin, but no stress fibers (Fig. 1F, left). Such puncta might normally merge in a cell-cell junction but have limited mobility at these hybrid interfaces (38).

The lack of clear stress fibers suggested a possible lack of focal adhesions—integrin-mediated structures that couple the cytoskeleton to the ECM around cells. This would make sense as there should be nothing for integrin binding to couple on an E-cadherin substrate, so we tested this by searching for expression of paxillin—a key protein in focal adhesions—in a mosaic culture. Here, we mixed a small percentage of Paxillin-GFP (green) MDCK cells with an Ecad:dsRed (red) population and demonstrated that cells straddling the Janus border formed focal adhesions only on the ECM side of the border, with apparent complete exclusion on the Ecad:Fc surface, emphasizing the complete biological and biophysical difference in cell attachment across the boundary (Fig. 1F, right; only the Pax-GFP cell is shown for clarity).

Finally, in addition to its other functions E-cadherin at lateral, cell-cell junctions help to separate the basal (bottom) and apical (top) surfaces of the cell and establish polarity in epithelia, so what happens to polarity if E-cadherin is instead presented to the basal surface? We tested this by staining a Janus tissue for podocalyxin—a key protein that marks the top of kidney cells (MDCK) and is related to microvilli and water transport. While we expected apical-basal polarity disruption, there was no apparent change across the Janus border, suggesting that cellular E-cadherin at the lateral junctions was sufficient for proper apical/basal separation (Fig. 1G).

### Cell-mimetic surfaces shut down collective cell migration and mechanical coordination

The Ecad:Fc side of the Janus substrate is akin to myriad ‘hands’ sticking out of the floor and attaching to cells through cadherin ‘handshakes’ rather than via focal adhesion ‘feet’ (hence the lack of Paxillin and actin stress fibers in Fig. 1F). This lack suggested that cells on the Ecad:Fc side of the Janus substrate cannot pull on the substrate via classical mechanobiological means, and might exhibit impaired migration. We investigated this with 60 hr time-lapse microscopy of large tissues grown across the Janus surface (Fig. 2A, Movie S2; SI Appendix). Using Particle-Image-Velocimetry (PIV), we were able to detect and quantify collective migration behaviors and observe powerful migratory suppression on the cell-mimetic side of the substrate (Figs. 2B, C). While cells on collagen ECM exhibited typical epithelial migration dynamics including the onset of contact inhibition at ∼40 hrs, cells on the Ecad:Fc side remained nearly 6X slower than their counterparts for much of the experimental time.

**Fig. 2.**
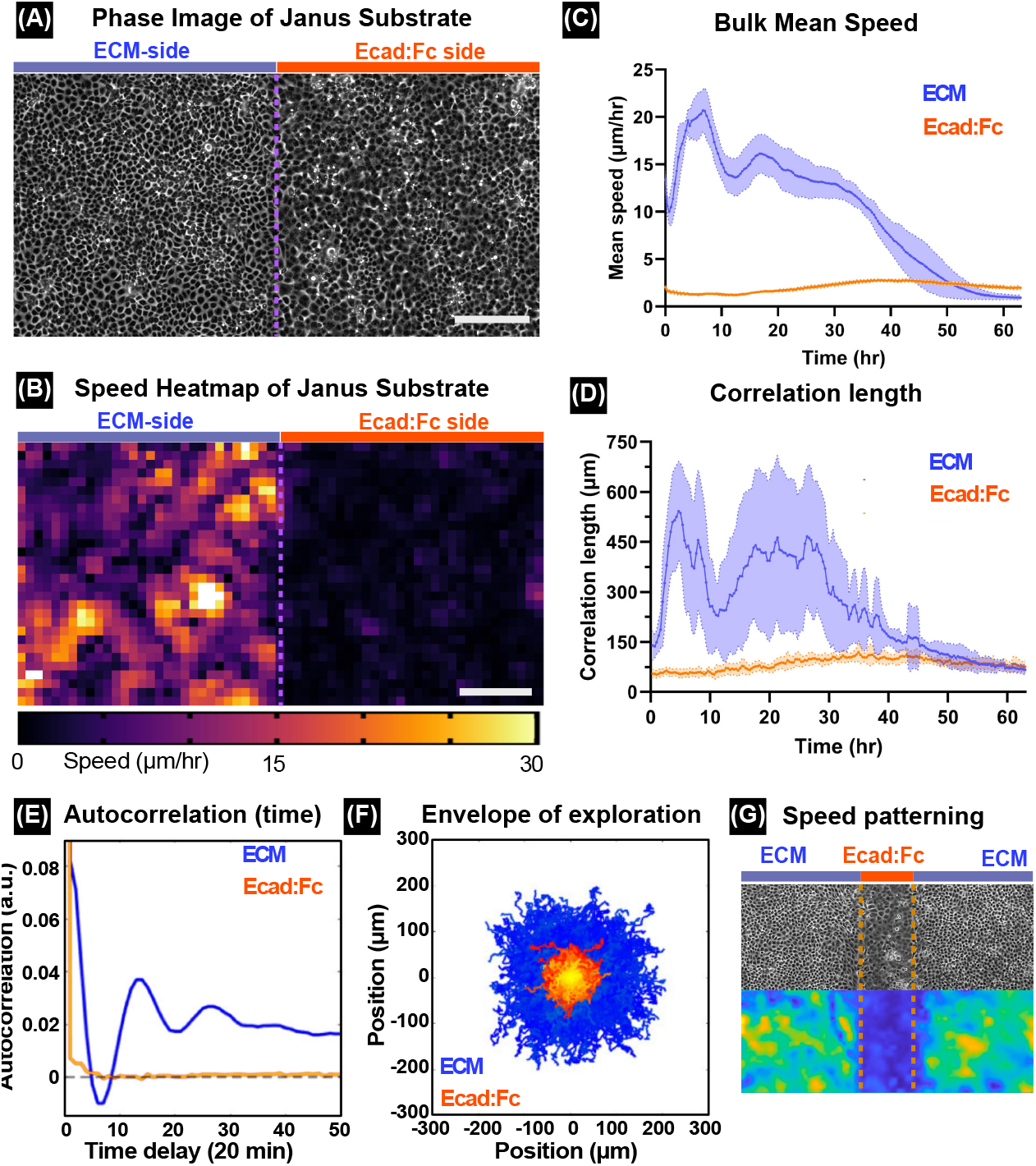
Ecad:Fc substrate locally suppresses collective cell dynamics. (A) Representative phase contrast image of single MDCK tissue seeded across a Janus substrate. (B) Representative speed heat map of single tissue on the Janus substrate. Scale bars: 250 μm. (C) Change of mean speed over time. (D) Correlation length comparison for cells within the ECM and Ecad:Fc regions of the Janus substrate (see Methods). (E) Velocity autocorrelation dynamics showing temporally periodic motion in regions of the tissue on ECM substrate, but non-persistent motion on Ecad:Fc subtrate. (F) ‘Hairball’ plot of cell tracks on ECM (blue) and Ecad:Fc (yellow-red) substrate, centered around the origin. n=16 across 3 experiments. (G) Phase contrast image and heat map of locally speed patterned tissue, with a narrow stripe of Ecad:Fc between ECM on either side. Ecad:Fc strip zone is 350μm wide; see Movie S3.

Reduced speed does not imply altered collectivity, so we next measured cellular crowd correlation dynamics. First, we determined the spatial correlation length of cells on the respective sides of the Janus substrate, which reflects the average domain size (neighborhood) where cellular directions are coordinated. Epithelia are well-known for long-range correlated migration on ECM substrates, and we measured coordinated neighborhoods of ∼10 cell diameters (450 μm). However, Ecad:Fc substrates reduced the correlated neighborhood size to only ∼1.5 cells (75 μm). This clearly demonstrates a loss of coordinated behaviors in those regions of the tissues, implying a disrupted flow of mechanical information between cells (Fig. 2D).

What were the cells doing if they weren’t migrating collectively? We measured this by tracking every cell and quantifying the temporal autocorrelation—a metric reporting how likely a given cell is to continue moving in the same direction it was previously going (Fig. 2E). Temporal autocorrelation*>*0 means correlation with past behaviors, while *<*0 indicates anti-correlation (e.g. moving the opposite way), and 0 implies no correlation at the given time interval. We observed an oscillating curve for cells on the ECM-substrate that never converged to 0; characteristic of the to-and-fro swirling patterns in epithelia (Movie S2). However, cells on Ecad:Fc exhibited no correlation with their prior motion. We visualized this difference with a hairball that overlays every measured cell trajectory in these samples showing that cell trajectories from the ECM-side (Fig. 2F, blue) enclose an envelope of nearly 10X the area as cells over the cell-mimetic, Ecad:Fc substrate (Fig. 2F, red/yellow region). We further emphasize the directedness of cells on ECM and the near diffusiveness of cells on Ecad:Fc using mean-squared-displacement analysis (Fig. S4). Integrating these data, cells on E-cadherin substrates not only lose their ability to mechanically coordinate with their neighbors, but they also lose their mechanical memory of past migration and become ‘migrastatic’ (39). Interestingly, this means that collective dynamics can be locally programmed into a single tissue by changing the substrate, which we demonstrated by patterning a narrow stripe of Ecad:Fc between regions of collagen ECM, which produced a small zone of static cells within the tissue (Fig. 2G; Movie S3).

### Mechanobiological information does not cross the Janus substrate boundary

What happens to collective cell dynamics near the Janus surface boundary where the highly collective motion on the ECM-side must transition to the near stationary behavior on the cell-mimetic side, i.e. how does information propagate? Epithelia are well known to transmit mechanical information over long distances via mechanical strain waves propagating through cell-cell tension/compression and transduced by cadherin junctions (40, 41), so we first analyzed the spatial velocity patterns far from and across the Janus boundary. Averaging PIV speed measurements revealed that the boundary not only induced a sharp drop in cell speed from the ECM-side to the Ecad:Fc-side, but that the reduction in speed occurred entirely on the ECM-side without any apparent change to cell speed on the Ecad:Fc side (Fig. 3A). For there to be no change in cell speed immediately to the right of the Janus boundary, mechanical information must not be propagating, so we calculated the actual strain waves (40–43) propagating across the tissue (Fig. 3B; Movie S4; SI Appendix), where ‘red’ means compression and ‘green’ means tension. These strain wave data revealed the expected highly correlated strain domains on the ECM-side and minimally correlated strains on the Ecad:Fc side. Unexpectedly, when we used a kymograph to plot the dynamics of these strain waves over time, we could see that strain waves traveling through the ECM-side of the tissue appeared to ‘reflect’ off the cells on the Ecad:Fc side of the Janus boundary rather than propagating across (Fig. 3C). To determine if this was true reflection, as would be expected from such waves hitting a mechanical barrier next to a tissue, we calculated the frequency power spectrum of strain waves moving towards and reflecting off of the Janus boundary. To ensure that strain waves were truly being reflected across the boundary, we sought to analyze any periodicity within the propagation of strain rate; our logic being that pure reflection will exhibit a singular characteristic frequency, whereas if waves over the ECM substrate were convolved with waves coming from cells over the Ecad:Fc substrate, our analysis would expose two (or more) characteristic frequencies. The temporal frequency with which these strain rate waves propagated through the tissue was calculated by performing Fourier transforms on signal traces (i.e., individual columns) within the strain rate kymographs (Fig. 3D, SI Appendix). By plotting the power spectra of strain rate propagation perpendicular to the boundary, one can see that there is a clear peak in the ECM spectra, representing a resonant frequency with which strain rate waves propagate within the tissue (Fig. 3E). Conversely, the power spectra on the Ecad:Fc substrate shows a canonical power cascade (45, 46), with no characteristic frequency. As such, we describe the boundary between the two substrates as a ‘pseudo-mechanical’ boundary, with properties such as strain wave reflection. As a final demonstration that the Janus substrate acts as an effective mechanical boundary between the two halves of the tissue, we calculated similar strain wave data where the tissue made contact with the silicone stencil walls on either side (Fig 3E, mid; see also Fig. 1A). Overlaying these data (Fig. 3E, bottom) shows similar periodicity in the reflection at a genuine tissue-mechanical wall interface to that at the Janus boundary where cells immediately on the Ecad:Fc side of the boundary appear to reflect mechanical information back into either side of the boundary rather than continuing to transmit the wave across the boundary.

**Fig. 3.**
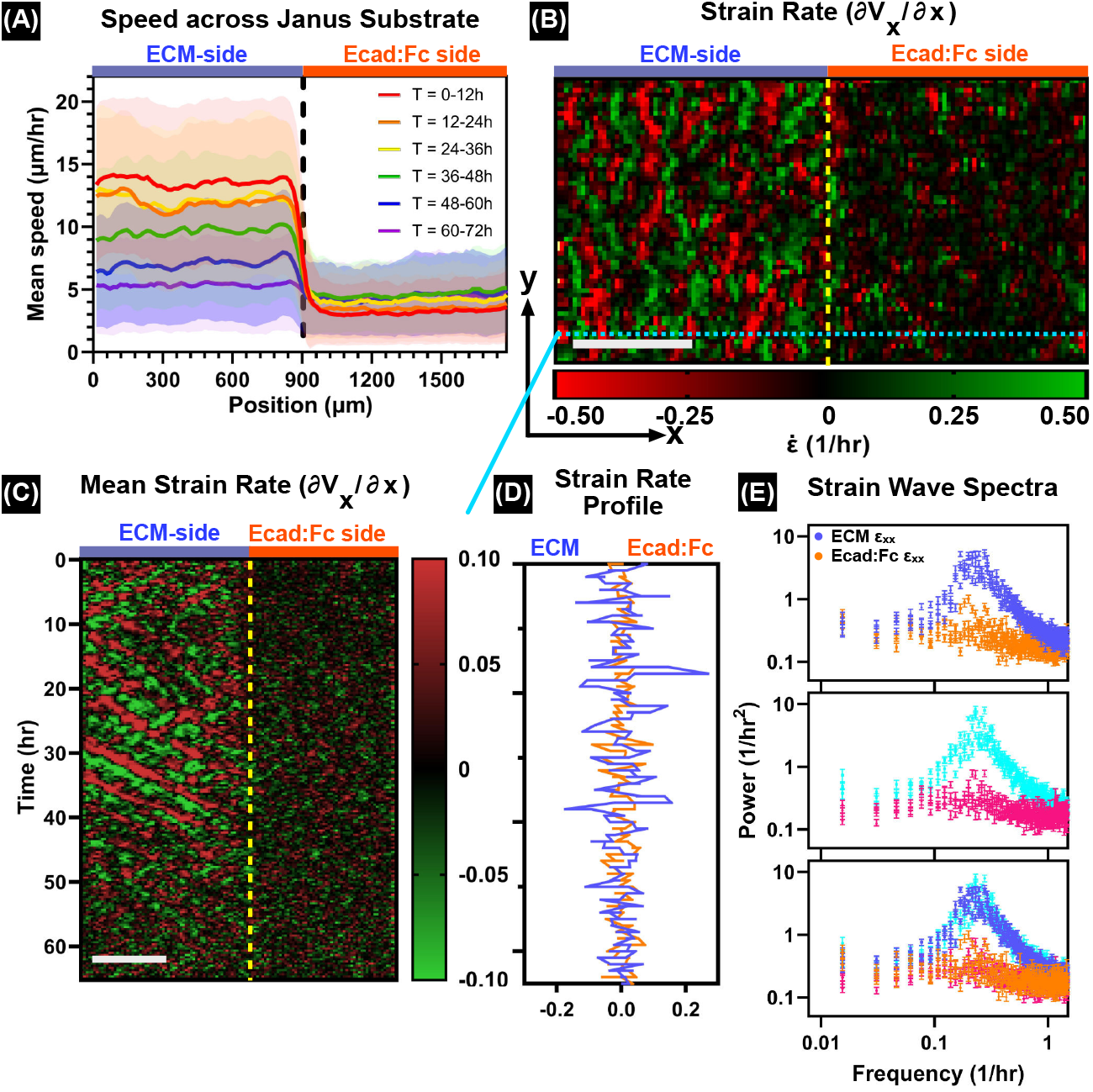
A ‘pseudo-mechanical’ boundary on the Janus substrate. (A) Spatial profiles of mean speed across the Janus substrate, binned into 12 hr periods. (B) Strain rate heat maps expose regions of highly correlated strain on the ECM substrate, but weaker, disordered regions on the Ecad:Fc substrate. (C) Strain rate kymographs exhibit reflection at the Janus boundary for ∼40 hr until contact inhibition. Scale bar: 50 μm (D) Cross sections of the strain rate kymograph show large-magnitude periodicity on the ECM substrate. (E) Power spectra of 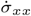 wave propagation on the ECM and Ecad:Fc substrate show resonance and a cascade, respectively. (top) Power spectra of 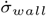 showing reflection at the tissue-PDMS boundary (middle). Overlay of 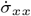 and 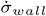 spectra, showing similar resonances between the two different boundaries (bottom). n=16 across 3 experiments.

### Reduced collective cell dynamics on cell mimetic substrates are not due to jamming

The typical explanation for reduced collective migration in tissue is ‘contact inhibition’, where dense cell packing in mature epithelia constrains cell rearrangements and divisions (47). We tested if this could explain behavior on the Ecad:Fc surfaces by measuring cell arrangement (Fig. 4A) and density on either side of the Janus substrate over time (Fig. 4B, left) based on fluorescent nuclei imaging (SI Appendix). Surprisingly, we found that average cell density increased more quickly on the ECM side (∼1.6X more cells were on the ECM side of the substrate than the Ecad:Fc side by the final time point), which agrees with density observations of keratinocytes on Ecad:Fc (20). Hence, increased cell crowding itself was not responsible for the damped behaviors on the Ecad:Fc side.

**Fig. 4.**
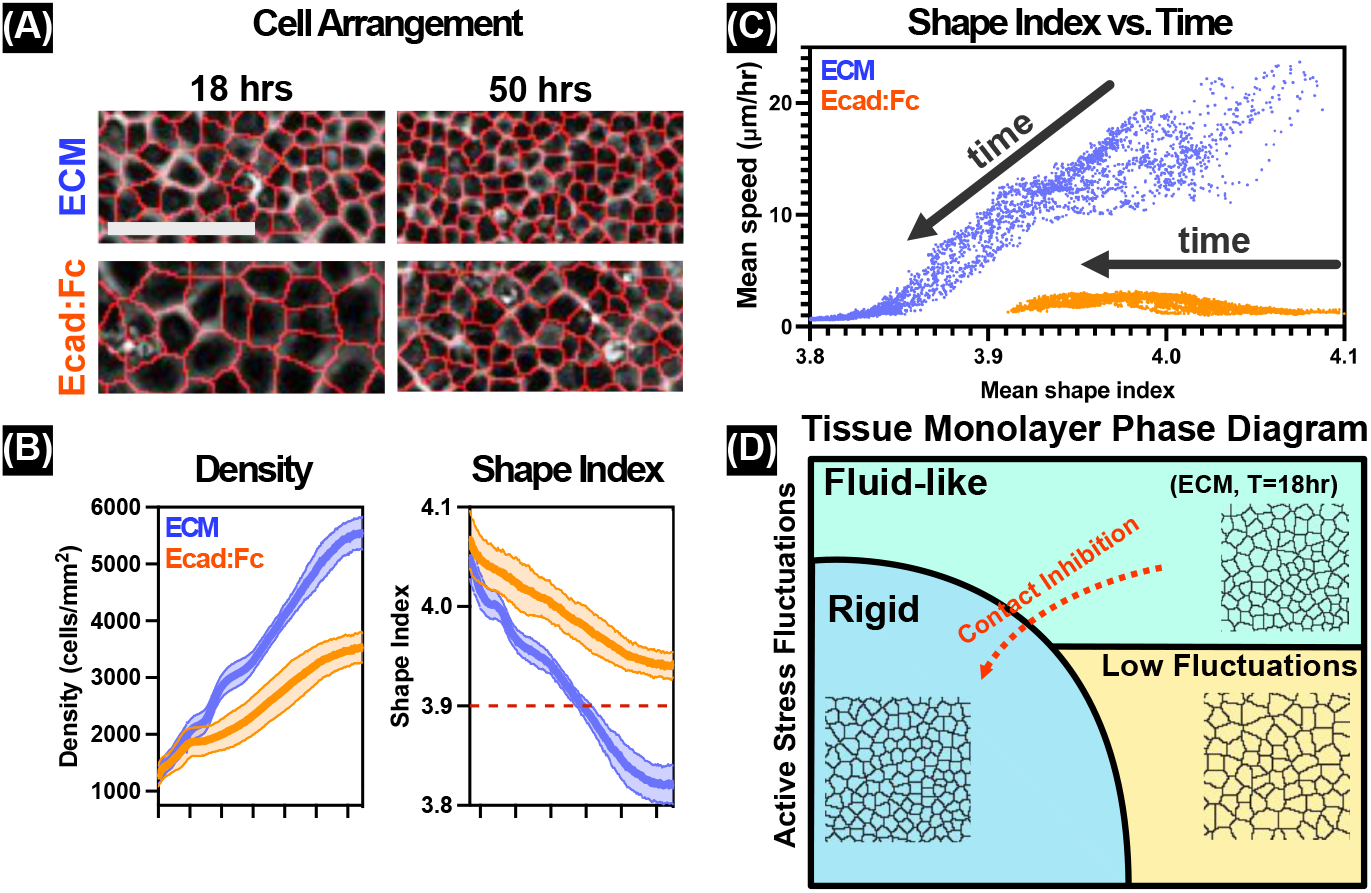
Ecad:Fc substrates breaks correlation between collective cell migration and the cell shape index. (A) Overlay of cell boundary mask and phase image, Scale bar: 100 μm). (B) Density (left) and cell shape index (right) evolution on two substrates. The dotted line indicates the threshold shape index, 3.9, for MDCK fluidity (44). n=16 across 3 experiments. (C) Mean frame speed versus Mean frame shape index plot. Arrows: time progression. (D) Tissue monolayer phase diagram. Tissue on ECM experiences contact inhibition over time (arrow). Tissue on Ecad:Fc stays in low fluctuation phase overall.

To better assess tissue mechanics, we evaluated the popular shape index metric that can predict if tissue has more solidlike or fluid-like characteristics (34); calculated as follows:

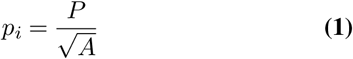

where *P* is the perimeter and *A* is the area of cell *i*. In MDCK epithelia it is assumed that a shape index *<* 3.9 implies that the tissue is more solid-like and likely to be jammed due to cortical tension dominating cell-cell adhesions, while a shape index *>* 3.9 suggests a more fluid-like tissue capable of larger fluctuations (34, 44). We calculated shape indices via Voronoi tessellation of fluorescent nuclear images (48, 49) for all cells over time (Fig. 4B, right). While the shape index of cells on the ECM-side dropped below 3.9 at ∼40 hrs as expected (contact inhibition) (50) (Fig. 2C), the shape index for tissues on the Ecad:Fc surface remained above 3.9, implying that near-stationary tissue on Ecad:Fc does not match the ‘solid-like’ model.

Directly comparing the mean shape index vs. mean speed (Fig. 4C) further emphasizes the different behavioral regimes between the two sides of the Janus surface. The ECM-surface drove a monotonic shift from high speed/high shape index to low speed/low shape index. By contrast, cells on the Ecad:Fc surface exhibited a near-constant speed over time accompanied by a weaker drop in the shape index. This behavior is not consistent with a simple solid-like/liquid-like binary classification of tissue rigidity. Recent work offers a different way to understand the effects of the Ecad:Fc surfaces on collective dynamics (51). All solid/fluid tissue models try to relate tissue rigidity to active stress fluctuations. Here, rigidity results from the stiffness of the monolayer tissue, while fluctuating active stresses within and between cells (contractility, traction forces, mitotic pressure) drive the characteristic, highly correlated motion patterns of epithelial cells on ECM substrates (47, 50). Contact inhibition describes how increased cell density drives rigidity and a transition to more solid-like behavior (e.g. the portions of our tissues on the ECM substrate side). For a tissue to slow down without an effective change to rigidity, the active fluctuations must be reduced, which would reduce effective agitation within the tissue(51– 53). This ‘low fluctuation phase’ has been hypothesized to result from disrupted focal adhesions (poor cell-substrate force generation) or reduced cell cycling, and would not require a shape index to drop below the critical value to result in an effectively immobilized tissue (51). Our data suggest that cell-mimetic substrates may be driving adhered tissues to such a low fluctuation state, which we further analyzed by explicitly studying both cell cycling and focal adhesion behaviors (Fig. 4D).

### Cell-mimetic surfaces reduce cell division frequency by increasing the G0/G1 phase of the cell cycle

Cell proliferation in the epithelium is a natural process to fill space and reach a homeostatic density and is strongly connected to collective cell migration (54) and mechanobiology. Our density data and that of others (55) suggested that cell-mimetic substrates may inhibit proliferation (Fig. 4B, left). We directly assessed this by using MDCK cells carrying the FUCCI cell-cycle marker where cdt1 is fused to mKO2 (magenta here) and geminin is fused to mAG1 (green here) (56). Hence, magenta fluorescence is proportional to progress through the G0/G1 phase of the cell cycle and green reflects S-G2-M. Post-mitotic cells are fully dark.

We adapted our previous methods of spatiotemporally mapping the cell cycle in large epithelia(50) to measure FUCCI dynamics on either ECM collagen surfaces or Ecad:Fc surfaces. By 15 hrs, we already observed striking asynchrony on either side of the Janus boundary with microscopy, indicating that cells cycled frequently on the ECM side of the interface but quickly locked into G0/G1 (magenta) on the Ecad:Fc substrate (Fig. 5A, Movie S5). Further, this asynchrony could be precisely patterned, which we demonstrated by patterning a thin band of Ecad:Fc within a collagen ECM backdrop. Fig. 5B and Movie S6 show how this resulted in a local region of high G0/G1 signal coinciding with the Ecad:Fc zone.

**Fig. 5.**
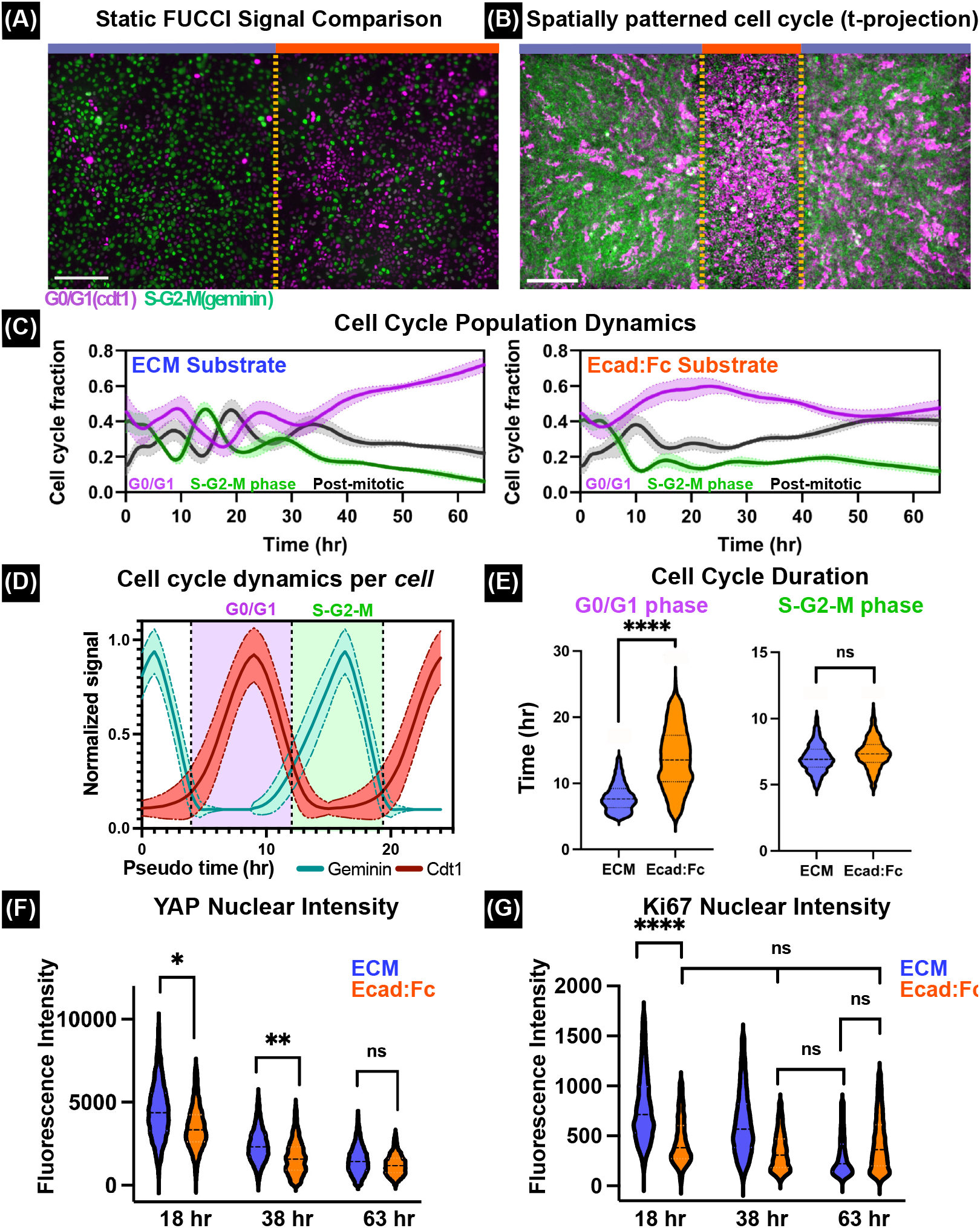
Ecad:Fc surfaces locally prolong the cell cycle. (A) Fluorescence imaging of MDCK cells reporting the FUCCI cell-cycle marker, seeded on the Janus boundary. The marker follows the geminin (green) and cdt1 (magenta) expression. Scale bar is 200μm. (B) Spatial patterning of the cell cycle by functionalizing a narrow stripe of Ecad:Fc between ECM on either side. The image shows a t-projection of 33 hr fluorescence time-lapse. Scale bar is 250μm. (C) Time evolution of cell cycle fraction on ECM (left) and Ecad:Fc (right); n=16 across 3 experiments. (D) Schematic of cell cycle duration quantification. The dotted lines indicate the threshold of G0/G1 and S-G2-M by measuring the time between the intersections of geminin and Cdt1 signals. (E) The cell cycle duration of G0/G1 phase (left) and S-G2-M phase (right) was measured during the initial 28 hr; n=24 across 6 experiments (1362 tracks on ECM, 1665 tracks on Ecad:Fc). (F) Quantification of YAP nuclear fluorescent intensity at selected time points; n=4 tissues with the following cell counts on ECM/Ecad:Fc. 18 hr: 5400/3800. 38 hrs: 8000/5200. 63 hrs: 13000/6600. (G) Quantification of Ki67 nuclear fluorescent intensity at selected time points. n=4 samples with the following cell counts on ECM/Ecad:Fc. 18 hrs: 5200/4170. 38 hrs: 7800/4300. 63 hrs: 11100/4200. ns, not significant, *P*<*0.05, **P*<*0.01, ****P*<*0.0001

To quantify population-level cell cycle regulation, we first measured the total population fraction of cells in G0/G1, S-G2-M, and post-mitosis over time (Fig. 5C) for either side of the Janus interface. Cells on ECM substrates exhibited stereotypical, out-of-phase cycling of cdt1/geminin (Fig. 5C, left), suggesting synchronized cell division up until 40 hrs where contact inhibition suppressed cycling (50). Cells on Ecad:Fc substrates behaved quite differently, with a total G0/G1 fraction consistently and significantly higher than the S-G2-M phase and a nearly 2X increase in post-mitotic cells (Fig. 5C, right).

We next performed cell cycle tracking on a single-cell basis for every cell over the entire experiment (SI Appendix), which allowed us to measure the absolute durations of the G0/G1 and S-G2-M phases (Fig. 5D) for cells on either side of the Janus boundary. Interestingly, Ecad:Fc substrates increased G0/G1 duration by nearly 2X (+73%) relative to cells on ECM substrates, but both substrate conditions had identical S-G2-M phases (Fig. 5E).

### Cell-mimetic, Ecad:Fc substrates delay the cycle primarily by inhibiting integrin signaling

Why would Ecad:Fc substrates increase the duration of the G0/G1 phase of the cell cycle? Cell-cycle regulation in dense epithelia is typically related to contact inhibition of proliferation through *β*-catenin/Wnt signaling and the Hippo/YAP pathway, and E-cadherin has been shown to play a role in both cases(55, 57–59), so we began our investigations here. Wnt activation allows the proper progression at the G1/S checkpoint by driving translocation of cytoplasmic *β*-catenin to the nucleus where it drives c-myc transcription (58, 59). As *β*-catenin also stabilizes E-cadherin, there is a potential competition between these processes and we hypothesized that hybrid junctions formed at the Ecad:Fc surfaces might deplete available *β*-catenin. However, our Western blots against E-cadherin and *β*-catenin (Fig. 1) showed no appreciable change across the Janus substrate, making sequestration less likely. We more directly tested the role of Wnt with an agonist (activator; CHIR 99021) to force Wnt activity; known to accelerate G1 (57, 58). To ensure Wnt was activated, we used TOPdGFP MDCK cells that report on *β*-catenin-induced transcription and increased fluorescence with Wnt activation (Fig. S5, left and middle). While Wnt activation accelerated G0/G1 times for both substrates, it did not close the gap in G0/G1 duration across the Janus interface (Fig. S5, right).

We next measured assessed Hippo/YAP signaling where YAP, a key cell-cycle regulator, should accumulate in the nucleus to promote cell cycle re-entry (59, 60). We observed a marked and sustained reduction in nuclear YAP levels at key time points for regions of tissue on Ecad:Fc surfaces vs. those on ECM surfaces (Fig. 5F, Fig. S6), suggesting that Ecad:Fc might delay cell cycle re-entry and that the G0-G1 transition might be affected. We tested this by measuring the distribution of nuclear Ki-67 signal from immunofluorescence (Fig. 5G, Fig. S7) as Ki67 is widely used as an indicator of actively cycling cells. Comparing Ki67 distributions in tissues on either ECM or Ecad:Fc surfaces at 18 hrs revealed significantly lower Ki67 levels on Ecad:Fc. Moreover, measured over time at 38 and 63 hrs, Ki67 levels on Ecad:Fc remained both low and self-similar. In fact, Ki67 behavior on Ecad:Fc substrates at all times most closely resembled Ki67 behavior in fully contact-inhibited tissues on ECM at 63 hrs (Fig. 5G, right). These data support the hypothesis that tissue attachment to a surface via E-cadherin may slow cell cycling by partially suppressing YAP and delaying the G0/G1 transition. However, while E-cadherin itself is known to modulate Hippo/YAP signaling via mechanobiology (55), it is interesting here that we did not detect altered E-cadherin (or catenin) protein levels across the Janus interface, which suggested to us that the something about the specific localization of cellular E-cadherin on Ecad:Fc substrates might be causing the changes to collective processes we observed.

One hypothesis that would connect altered collective migration and cell cycling relates to the pure mechanics of how cells migrate. A key regulator of crawling cell migration is integrin-mediated focal adhesions (‘feet’) (1, 3), and we had already established that Ecad:Fc surfaces appeared to exclude the formation of focal adhesions (Fig. 1F, right), so we next investigated the role of focal adhesions. Interestingly, integrin signaling has been shown to vary directly with YAP signaling, meaning that integrins do play a role in cell cycle variation (61, 62). To better study this, we performed a parameter sweep with tissues cultured on substrates functionalized with different direct blends of collagen/Ecad:Fc, effectively altering the relative abundance of ECM vs. cell-cell-adhesion ligands. Here, we premixed Col-IV and Protein A/G at different ratios and then functionalizes single glass surfaces with these blends (SI Appendix). As expected, higher proportions of ECM resulted in weaker basal Ecad:dsRed recruitment in cultured tissues and increased phosphorylation of focal-adhesion-kinase (pFAK; Figs. 6A,B, left). Total Internal Reflection Fluorescence (TIRF) imaging of paxillin itself, a key ‘foot’ protein, highlighted how the distribution of actual focal adhesions diminished with increasing Ecad:Fc proportions (Fig. 6A, right). More surprising were the effects on G1 duration, where we found that all of the blends we tested that contained any level of collagen exhibited statistically identical G1 transit times and it was only when we completely eliminated the ECM component from the blend (pure Ecad:Fc) that we observed the dramatic increase in G1 transit time (Fig. 6C). Hence, cell-mimetic Ecad:Fc surfaces appear to regulate the cell cycle via exclusion of focal adhesions and integrin signaling, rather than by more direct means.

**Fig. 6.**
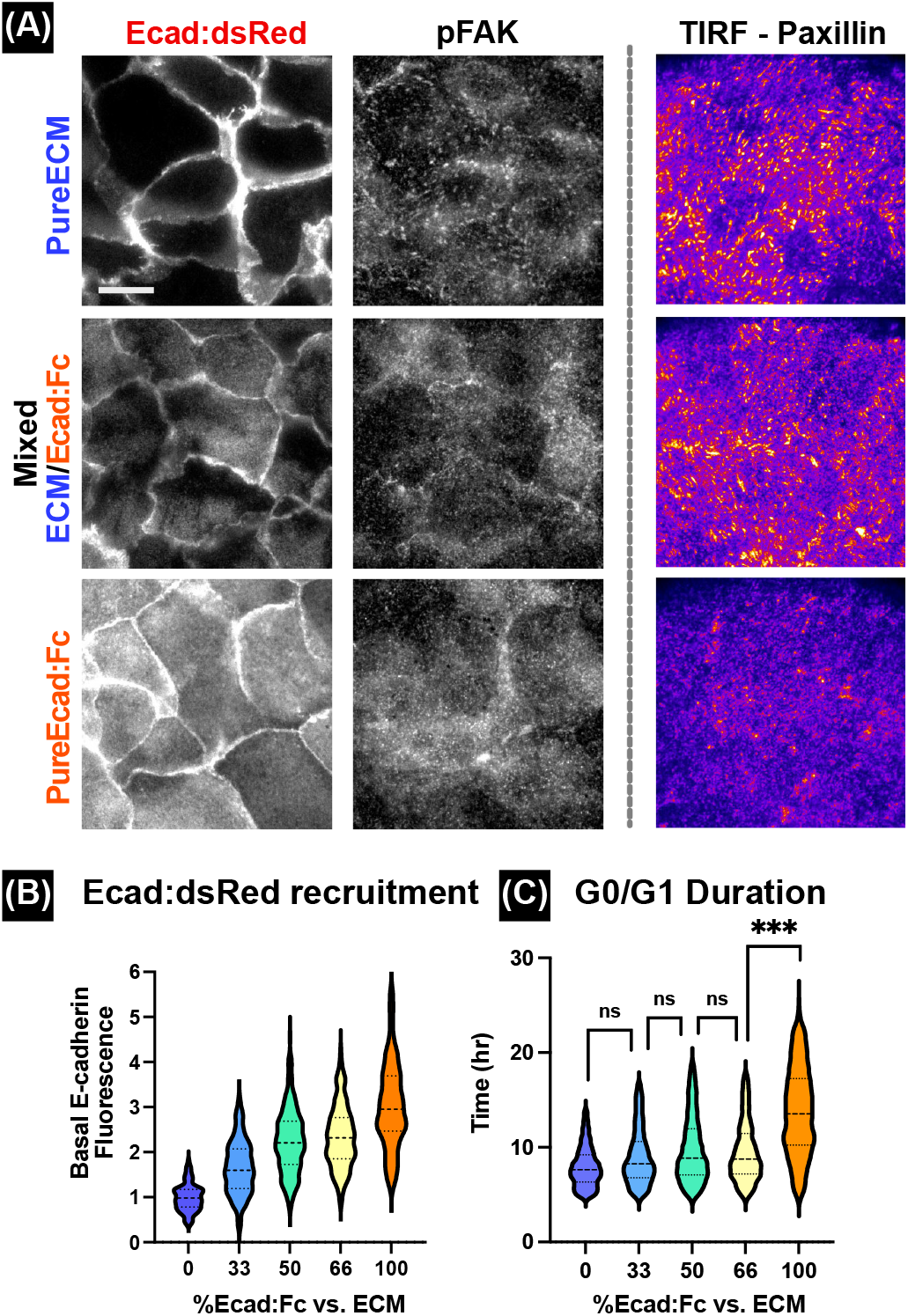
Ecad:Fc substrates exclude focal adhesions, which is correlated to cell cycling. (A) *top row:* Basal immunofluorescence imaging of Ecad:dsRed MDCK tissue on pure ECM; *middle row:* 50/50 mixed ECM/Ecad:Fc; and pure Ecad:Fc substrate (*bottom row*), showing Ecad:dsRed signal (*left column*), immunostaining of pFAK (*center column*), and TIRF imaging of paxillin (*right column*). Scale bar: 15 μm (B) Quantification of the normalized average basal Ecad:dsRed signal on mixed substrate with different Ecad:Fc/ECM concentration. 120 samples for 0, 100% Ecad:Fc substrate and 240 samples for 33, 50, 66% Ecad:Fc substrates across 3 experiments. (C) G0/G1 duration of cells on mixed substrate with different Ecad:Fc/ECM concentration. 1362,1665 tracks for 0, 100% Ecad:Fc substrates across five experiments. 1392, 1672, 1611 tracks for 33, 50, 66% Ecad:Fc substrates across 6 experiments. ns, not significant, ***P*<*0.001

## Discussion

The Janus surface is a unique material that contains separated ‘substrate-like’ (ECM) and ‘cell-like’ (Ecad:Fc) cell adhesion regions allowing a single tissue to be grown across these different regions. We likened this interface to a ‘floor’ on the ECM side and a field of grasping ‘hands’ on the Ecad:Fc side, allowing the material to trigger entirely different biological and biophysical responses in contacting regions of the tissue, and allowing us to investigate how the manner in which cells attach to the substrate reprograms collective cell dynamics. The nearly complete exclusion of focal adhesions on the Ecad:Fc side in favor of strong E-cadherin adhesion on the cell-mimetic side of the interface (Figs. 1,6) highlighted the binary nature of the interface and validated the precision of the interface between the two sides.

The overall trend on collective behaviors that we observed was that the cell-mimetic, Ecad:Fc side of the interface appeared to promote some form of forced homeostasis. Mechanically, this manifested as a near complete cessation of collective migration, indicated both by the loss of spatial correlation across the tissue (Fig. 2D), as well as the loss of persistent migration at the individual cell level (Fig. 2E), which was molecularly supported by the disruption of actin stress fibers (Fig. 1F) and focal adhesions (Fig. 6A). Physiologically, we observed a dramatic slow-down in the G0/G1 phase of the cell cycle. Here, the extremes of the cell cycle distribution data are instructive. While the mean G0/G1 transit time for cells in tissues on Ecad:Fc increased from 8 hrs to ∼14 hrs, upper ends of the distribution showed some G0/G1 times of nearly 24 hrs (Fig. 5E).

The underlying basis for these phenomena were not attributable to the standard explanations of cellular crowding/density causing contact inhibition of proliferation and locomotion based on the shape index metric and treating the tissue as solid-like or fluid-like. We instead speculate that cell-mimetic surfaces induce a previously postulated low-fluctuation state (51). However, the overall effects of the cell-mimetic substrate—migrastasis and reduced proliferation—clearly resembled the broad phenotype of contact inhibition.

These data highlight that how cells attach to a substrate matters more than the physical act of attachment. For instance, the portion of tissue on the ECM side of the Janus substrate often visually resembled the portion of tissue on the Ecad:Fc side and both formed intact epithelia, but with obvious behavioral differences. Moreover, the fact that Western blotted total E-cadherin and *β*-catenin protein levels did not differ in tissues on the two substrates (Fig. 1E), along with the data on blended/mixed substrates (Fig. 6B, right), emphasize that the exclusion of focal adhesions from the basal surface of the cell, played a major role in many of the effects we observed. Finally, throughout this work we demonstrated that the interface between the two sides of the Janus surface created remarkably sharp separation of function and behavior even at the level of single cells straddling the boundary (Fig. 1F). In particular, we found that mechanical information from one side of the tissue was unable to propagate across the Janus interface, instead reflecting akin to what happened when a tissue encountered a true, inert physical barrier (Fig. 3E). This sharp separation allowed us to spatially pattern both collective migration within a larger tissue (Fig. 2G), and even create sharp zones of altered cell cycling (Fig. 5B). These findings highlight the potential versatility of Janus biomaterials to inducing complex behavioral differences that may benefit future biomedical implants, such as by stabilizing the behavior of soft tissues at hard implant interfaces to reduce tissue separation and infection risks in percutaneous implants, for example (21, 22).

## ACKNOWLEDGEMENTS

Support for this work was provided in part by NIH Award R35 GM133574-03 (D.J.C.). We also thank members of CohenLab for advice and support.

