## Supplementary_Information for "E-cadherin biointerfaces reprogram collective cell migration and cell cycling by forcing homeostatic conditions"

### Supporting Information Text

#### Materials and Methods.

**Cell maintenance.** All experiments were performed with MDCK-II cells of the following types: stable lines expressing Ecad:dsRed, FUCCI, or TopDGFP; and a transfected mosaic sample with Paxillin-GFP. All MDCK cell lines were cultured in low-glucose/low-bicarbonate Dulbecco's Modified Eagle's Medium media with phenol red (D5523-10L, Sigma), supplemented with 1g/L sodium bicarbonate (S5761-500G, Sigma), 10 % (vol/vol) fetal bovine serum (S11550, Atlanta Biologicals), and 1 % (vol/vol) streptomycin/penicillin (15140-122, Gibco). Cell culture media was exchanged every two days unless otherwise noted. Cells were maintained at 37 ℃ and 5 % CO2 in humidified air. All cells tested negative for mycoplasma. (CUL001B, MycoProbe)

**Janus substrate functionalization.** A glass bottom dish (CellVis) was first hydroxylated with the plasma cleaner (PDC-001-HP, Harrick Plasma) at 50W excitationfor 3 minutes under low pressure (<300 mTorr). Other than the use of glass, the functionalization chemistry followed Cohen et al., PNAS 2016 (1). Plasma-treated glass bottom dishes were immediately immersed with the mixture of 2 % (vol/vol) triethoxysilylundecanal (aldehyde reactive group; SIT8194.0, Gelest), 2 % (vol/vol) triethylamine (T0886, Sigma) in pure ethanol (459844-500ML, Sigma) for one hour at room temperature. Dishes were washed three times with pure ethanol and baked at 80 °C for 3 hours to complete the condensation reaction.

A silicone stencil of 250 µm thickness (Bisco HT-6240, Stockwell Elastomers) was cut to the desired geometry using a Silhouette Cameo vinyl cutter and transferred to the silanized substrate as a mask. We carefully designed the stencil to initially cover the region intended for ECM coating, which we only removed after Ecad:Fc functionalization (Fig. 1A, for example). We first functionalized the initially exposed area with Protein A/G(PAG; 21186, Thermoscientific; 200 µg/ml) diluted in cyanoborohydride coupling buffer (C4187-500ML, Sigma) to improve coupling to the aldehyde silane for one hour at 37 ℃ or overnight at 4 ℃. Microwells within the stencil were washed five times with PBS with 1 mM CaCl2 (C34006, PromoCell). After Protein A/G, we added purified E-cad:Fc (Biolegend, 779906; 20 µg/ml in our Ca-supplemented PBS). Ecad:Fc working solution was added to the microwell and incubated at 37 °C for 3 hours, after which the dish was washed with Ca-PBS and then blocked with Bovine Serum Albumin in Ca-PBS (BSA; A1595-50ML, Sigma; 0.1 %) for 1 hr at room temperature. This blocking step was critical to prevent non-specific binding of serum and ECM proteins in the following steps. After blocking, we removed the trapdoor region of the stencil with a scalpel (Fig. 1A) and backfilled the dish with collagen IV solution (C7521-5MG, Sigma; 50 ug/ml) at 37 ℃ for 30 minutes, after which we washed three times with Ca-PBS. For the fluorescence validation, GFP fluorescent gelatin was mixed with collagen IV at 1:5.

To functionalize the mixed substrates with different Ecad:Fc/ECM concentration, PAG was directly mixed with Col IV in different ratio and incubated at 37 °C for 1 hour. After washing three times with Ca-PBS, Ecad:Fc was incubated.

**ELISA.** The glass bottom 96 multiwell plates were first functionalized with Ecad:Fc as described above. The concentration of the Ecad:Fc solution was adjusted with Ca-PBS. Non-specific binding to the glass and detection antibody Fc binding to PAG were prevented by blocking with the mixture of BSA (0.1x) and human Fc fragments (100 µg/ml) for 1 hour at room temperature. After washing the blocking buffers, we incubated the primary antibody against the extracellular domain of E-cadherin (16-3249-85, Invitrogen; 1:100) on the substrate for 1 hour at room temperature. After washing with the calcium PBS, the HRP-conjugated secondary antibody (31470, Invitrogen; 1:5000) was incubated for 1 hour at room temperature. The amount of functional E-cadherin on the substrate was detected with the o-phenylenediamine substrate (34005, Thermo Scientific; 1 mg/ml) by measuring the OD absorbance values at 450 nm with the plate reader (Infinite 200Pro, Tecan).

**Tissue patterning.** Cells were washed with PBS and detached from the tissue culture plastic dish by incubating the cells in TrypLE (12604-013, Gibco) at 37 ℃ for 7 minutes. Cell solution in TrypLE was diluted with the culture media and centrifuged at 1500 RPM for 3 minutes. The supernatant was aspirated out and the cell pellet was resuspended into the fresh culture media to acquire desired concentration (∼2.2E6 cells/mL). The suspended cell solution was well-triturated and seeded within the stencil on the functionalized substrate (2, 3). Seeded cells were allowed to adhere to substrates for 2.5 hrs in the incubator before the dish was flooded with DMEM. Humidity was maintained with a ring of media around the periphery of the dish).

**Microscopy.** Both phase contrast and epifluorescence images were acquired with an inverted Nikon Ti2 microscope using NIS Elements software and a Nikon Qi2 camera. Live-cell imaging was performed with environmental control that maintained 37 ℃ and humidified 5 % carbon dioxide. All time-lapse images were captured every 20 minutes for 63 hours. 4 X epifluorescence was performed with 12 % Lumencor SOLA intensity (TRITC cube) and 13 % (GFP cube), 400 ms exposure time, and 5.1x Qi2 gain. For higher magnification (20 X and 60X), imaging conditions were optimized for each filter to improve the signal-to-noise ratio while avoiding phototoxicity and photobleaching.

Confocal microscopy was performed at the Princeton University Confocal Imaging Core Nikon Center using a Nikon Ti2 inverted microscope with a Yokogawa W1 spinning disk using the laser lines with 405 nm, 488 nm, 561 nm, and 647 nm wavelengths. Images were captured using a 60x oil immersion objective with a Hamamatsu BT Fusion CMOS. Total Internal Reflection Fluorescence (TIRF) imaging was performed on a Nikon Ti-E with Nikon TIRF and a high NA 60X objective.

**Immunofluorescence Staining.** Tissues were washed twice with PBS and fixed with 4 % (vol/vol) paraformaldehyde solution (15710, Electron Microscopy Science) diluted in PBS for 15 minutes at room temperature. Tissues were washed five times with PBS and permeabilized with 0.1 % Triton-X-100 (T8787-100ML, Sigma) diluted in PBS for one hour at room temperature.

After washing three times with PBS, blocking solution (2 % BSA, 1 % normal donkey serum, 1 % goat serum, 50 mM ammonium chloride, 0.05 % sodium azide in PBS; filtered through 0.45 um filter) and pure human IgG Fc fragments (200 ug/mL; ab90285, Abcam) were mixed and incubated overnight at 4 °C or for 3 hours at room temperature to prevent the non-specific binding and Fc-PAG binding with antibodies, respectively.

Primary antibodies used for immunostaining were: Mouse anti-paxillin antibody (1:100; 610051, BD Biosciences), Mouse anti-podocalyxin (1:200; MABS1327, EMD Millipore), Alexa Fluor 488 phalloidin (1:1000; A12379, Invitrogen), Rabbit anti-phospho-FAK(Tyr397) (1:200, 700255, Invitrogen), Rabbit anti-Ki67 (1:200, ab16667, Abcam), Mouse anti-YAP (1:200, sc-101199, Santa Cruz).

Secondary antibodies used for immunostaining were: Alexa Fluor 647 goat-anti-mouse (1:500; A21235, Invitrogen), and Alexa Fluor 488 rabbit-anti-mouse (1:500; A11059, Invitrogen). Cy5 goat anti-rabbit (1:500; A10523, Invitrogen) All antibodies were diluted to a working concentration with blocking solution. For nuclear staining, one standard drop of NucBlue Reagent (R37605, Invitrogen) was added to the secondary antibody mixture.

**Migration analysis.** We used particle-image-velocimetry (PIV) to generate velocity vector fields for timelapse analyses using 4X phase contrast imaging and the PIVLab package for MATLAB (4). We used 2-pass FFT analysis using a box 1 size of 64 pixels and box 2 size of 32 pixels with 50% overlap.

The mean bulk speed of tissue was calculated by the arithmetic mean of the speed of the individual PIV boxes at each frame. Velocity profiles across the Janus substrate surface were calculated by averaging over columns of the PIV output. The resulting averaged velocity profiles were binned into 12 hr periods and their mean and standard deviation were plotted.

Velocity-velocity correlations were calculated using a custom MATLAB script utilizing cell tracks across the entirety of the experiment (generated via TrackMate in FIJI). Track data was used to calculate instantaneous velocities for every cell at every time point, and correlation between cells was calculated according to Cavagna et. al.(5). By subtracting the global mean, correlations were calculated on the residuals, meaning that the correlation length can be chosen to be where the correlation function crosses zero. Correlation lengths at each time point were then averaged across all cells in the bulk of each substrate and their mean and standard deviation was plotted.

**Strain Rate.** The strain rate tensor for each tissue was calculated as follows

$$\dot{\sigma} = \frac{1}{2}(\nabla V_x + \nabla V_y) \qquad [1]$$

Where v is the velocity field of the tissue at time t, calculated using PIV (see *Migration Analysis*). Strain rate heat maps were produced by plotting the first diagonal component ($\sigma_{xx}$) of the strain rate tensor at every position within the tissue. See (6).

To visualize strain rate wavefront dynamics, average kymographs were calculated as follows: a row m of the MxN strain rate heat map was transformed into a TxN kymograph (T is the number of time points in the experiment), where each row t is m(t). Each MxN strain rate heat map will result in M TxN kymographs, which are then entry-wise averaged.

**Spectral Analysis.** Power spectra were calculated by first performing a temporal fast Fourier transform on the columns of the aforementioned strain rate kymographs, which was physically representative of the temporal strain rate fluctuations in a particular point within the tissue. The fast Fourier transform was calculated using the built-in MATLAB function fft(), which calculates the discrete fast Fourier transform. Once calculated, we took the square of the Fourier transform to retrieve the power spectrum, which was then averaged over all columns in the strain rate kymograph. The mean and standard deviation were plotted on a logarithmic scale.

**Image analysis.** Z-stack images were acquired with confocal imaging (See *Microscopy*) and Z-projected by summing all stacks. In this z-summed image, basal E-cadherin recruitment at the tissue-substrate interface was quantified using the Measure tool within Fiji. 80 x 80 px squares were drawn in the middle of each cell to measure the average intensity of TRITC signals within the square. For junctional E-cadherin intensity quantification, we used the FIJI tool *Junction Mapper* to get the mask file of cell-cell junctions and measured the tissue-averaged intensities using Matlab regionprop function (7). For both quantifications, background signals were estimated by averaging the fluorescent intensities at the cell-free zone within the same dish. The signal was subtracted with the background signal and normalized by the average of intensities on ECM substrate.

For nuclear intensity quantification of YAP, Ki67, and TOPdGFP signal, Nuclear mask is segmented from nuclear staining image using StarDist imageJ plugin and eroded by 5 pixels (8). Mean intensity of fluorescent marker of each nuclei is measured via regionprop MATLAB function.

**Cell shape index.** Nuclear images of MDCK tissue were reconstructed from 4x phase images using our in-house Fluorescence Reconstruction Microscopy tool (9). Cell boundary images were obtained by Find Maxima (segmented particle) Fiji plugin. The area and perimeter of each cell were measured using the Analyze Particle function of Fiji. The shape index of each cell was calculated as follows (10): $\frac{Perimeter}{\sqrt{Area}}$

**Cell density.** Nuclei positions are detected by applying Find Maxima Fiji plugin to Nuclear images of MDCK tissue reconstructed from 4x phase images. The local density map was calculated based on the number of cells in a fixed ROI. The ROI was 32 pixels by 32 pixels square and was scanned over the image with a 50 % overlap. The total density of tissue was calculated as (total number of cells in the tissue/area of the tissue) in MATLAB.

**Cell cycle analysis.** Cell cycle fraction analysis was done with 4x phase, RFP, and GFP timelapse images. To normalize the histogram of RFP and GFP channel images, 'enhance contrast' and rolling-ball background subtraction were applied to the images in FIJI. Using a custom MATLAB (Mathworks) script, the mean RFP and GFP intensity of each nuclei were measured. Cell cycle stage of each cell was determined as described in (2) by comparing mean RFP and GFP intensity with manually chosen threshold values as a G0/G1 phase (RFP above threshold), S-G2-M phase(RFP below threshold and GFP above threshold), and post-mitotic phase (both RFP and GFP below the threshold). Local cell cycle fraction was calculated as the ratio of the number of cells in each cell cycle to the number of total cells in the density calculation box. (see *Cell density*)

For cell cycle duration analysis, RFP and GFP fluorescent intensities of individual nuclei were tracked from 3-channel image stacks (RFP, GFP, reconstructed nuclei) via Trackmate Fiji plugin. 20x timelapse (0-28 hours) images were used for this analysis. Further duration analysis was done by custom MATLAB scripts. Each channel signal of individual tracks was internally normalized by setting 0 for the lowest value and 1 for the highest value of the channel signal of the track. G0/G1 duration was defined as the timespan between intersection points of normalized RFP and GFP signal which has a higher normalized RFP signal than normalized GFP signal, and vice versa for S-G2-M duration.

FUCCI signal dynamics for each tracked cell were individually normalized and analyzed before compositing.Mean signal peaks were obtained by aligning the peak positions of each signal peak. Mean signal peak graphs of RFP and GFP channels were overlaid to reconstruct the FUCCI signal evolution plot. Phasing between RFP and GFP peaks was adjusted based on mean G1 duration and S-G2-M duration measured in cell cycle duration analysis. From individual signal peak data, rise and fall time was calculated by measuring time taken for the signal to rise or fall from/to 15 % and 85 % of peak signal. The entire process was done with a custom MATLAB script.

**Western blot.** To improve the Western blot signal strength, the entire glass bottom dish was functionalized with either Ecad:Fc or ECM, and 300 $\mu$L of MDCK cell solution was seeded on the entire glass bottom part without stencils. After allowing cells to adhere to substrates, media was added to fill up the dish. After 18 hours of incubation at 37C, two dishes per condition were transferred to an ice bucket and washed twice with ice-cold PBS. Tissues were collected by scraping the dish in 400 $\mu$L of NP-40 lysis buffer (BP-119, Boston Bioproduct) with protease inhibitor cocktail (10 $\mu$L per mL; AR1182, Boster) and lysed for 30 minutes on the rocker. The cell solution in lysate was centrifuged at 4 ℃ and the supernatant was aliquoted to be stored at - 80 ℃ for future uses.

Due to variations of cell density at 18 hours, the BCA assay kit (23227, Thermoscientific) was used to quantify the total protein concentration in the lysate solutions collected from two substrates. With the absorbance function on the plate reader, the standard curve was first generated with the BSA solution provided in the BCA kit. The amount of total protein in the lysate solution was quantified using the absorbance measurement and BSA standard curve. Sample volume in the gel loading solution was adjusted to match the protein concentration of the lysate from ECM substrate and EcadFc:substrate.

The gel loading solution was prepared by mixing the appropriate amount of lysate sample, Tris-Glycine SDS sample buffer (LC2676, Novex), NuPAGPE sample reducing agent (NP0004, Invitrogen), and deionized water. The mixture was heated at 85 ℃ for 2 minutes before being loaded to the gel. Proteins were separated with NovexTM WedgeWellTM 4-12 % gel (XP04122BOX, Invitrogen) with Mini Gel Tank (A25977, Thermofisher) in 1x diluted Tris-glycine SDS running buffer (BP-150, Boston BioProducts) at constant 225 V for 30 minutes (PowerEase 300W, Life Technology). The reference molecular weight was tracked with protein dual-color standards (1610374, Biorad).

After separation, proteins were dry transferred to the nitrocellulose membrane with the iBlot 2 (IB21001, Invitrogen) by placing the gel in between the iBlot 2 NC Mini Stacks (IB23002, Invitrogen). After the transfer, the membrane was incubated in Revert 700 total protein stain kits (926-11011, Li-cor) for western blot normalization and rinsed twice with Revert 700 wash solution. The membrane was imaged with LI-Cor Odyssey CLx imager using the appropriate channel to acquire total protein images. An example total protein stained gel image shown in Supplementary Figure 3. The total protein stains were then removed with Revert destaining solution in the kit to perform the blotting against E-cadherin and $\beta$-catenin.

In order to prevent the non-specific binding to the nitrocellulose membrane, the membrane was blocked with the 1X iBind FD solution (SLF1019, Invitrogen) at room temperature for at least 30 minutes. Primary and secondary antibodies were diluted to desired concentrations with 1X iBind FD solution and added with the iBind FD 10 % SDS to reduce the background signal. The iBind card (SLF1010, Invitrogen) was placed in the iBind Western device (SLF1000, Life Technology) with an appropriate amount of antibody solutions and 1X iBind FD solution. Antibody binding was then performed within the device by placing the device at room temperature for at least 2.5 hours. The membrane was washed with DI water twice and antibodies were detected with LI-Cor Odyssey CLx imager using 680 channel. Western blot protein quantification was performed via measuring the area of band and signal in Fiji. The signal was subtrated with background signal and normalized with the total protein stain signals. A rectangular ROI was selected to cover each lane and its integrated signal was calculated by (ROI area) X (ROI mean intensity - background mean intensity).

Antibodies used for Western blot staining were: Mouse Anti-E-cadherin C-terminal antibody (1:1000; 610181, Bd Biosciences), Mouse Anti-$\beta$-Catenin antibody (1:2000; 610153, Bd Biosciences), IRDye 680 RD goat anti-mouse (1:4000; 926-68070, Li-Cor).

**Statistical test.** Mann-Whitney U test is adopted to test statistical difference between two data sets. 50 observations are randomly sampled with replacement from each data set. P-value is calculated by running Mann-Whitney U test between the subsamples. Mean P-value of 50 iterations of this process is used for deciding significance.

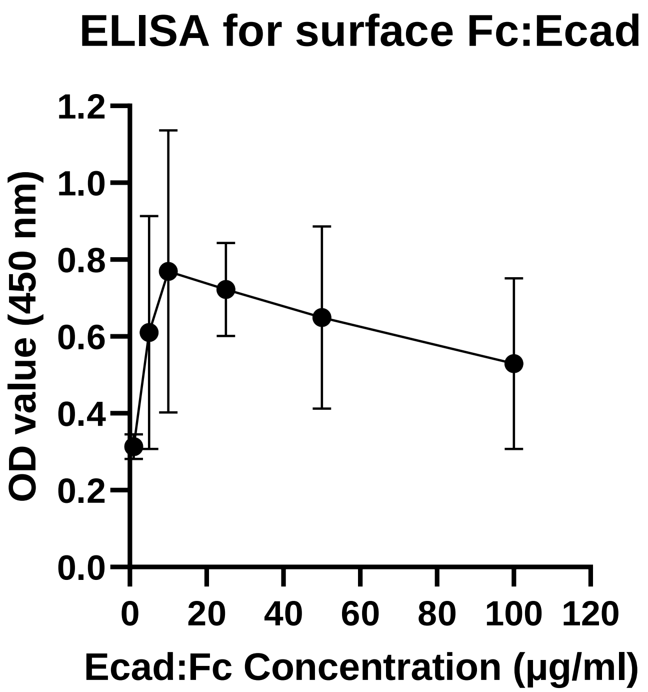


**Fig. S1.** Detection of the functionalized Ecad:Fc on the glass substrate via ELISA assay. The maximum OD value was read around 10 $\mu$g/ml, so we used 20 $\mu$g/ml Ecad:Fc concentration for our study. (n=4 for each condition)

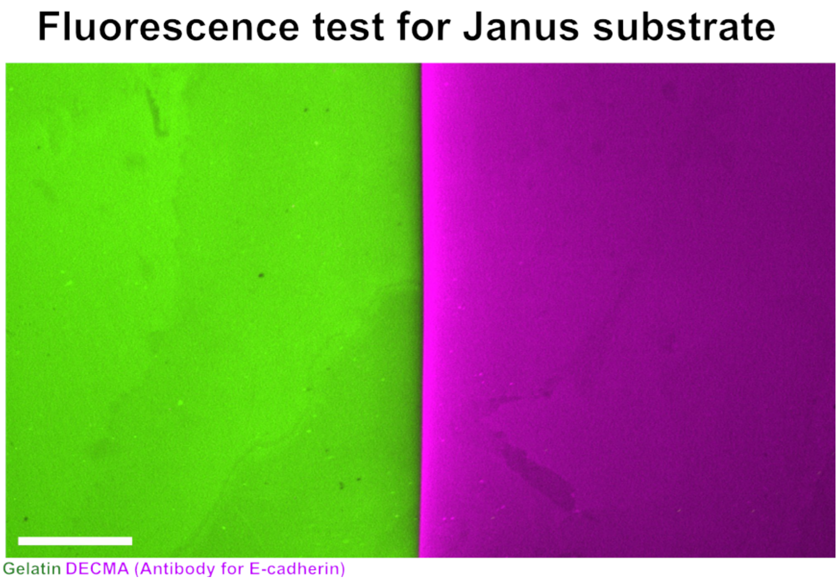


**Fig. S2.** Validation of Janus substrate functionalization, showing no bleed-through over boundary in fluorescence image. After Ecad:Fc functionalization, Collagen IV mixed with fluorescence gelatin (80/20 vol/vol %) was backfilled, followed by the immunostaining against surface E-cadherin using DECMA antibody. (Scale bar: 200 $\mu$m)

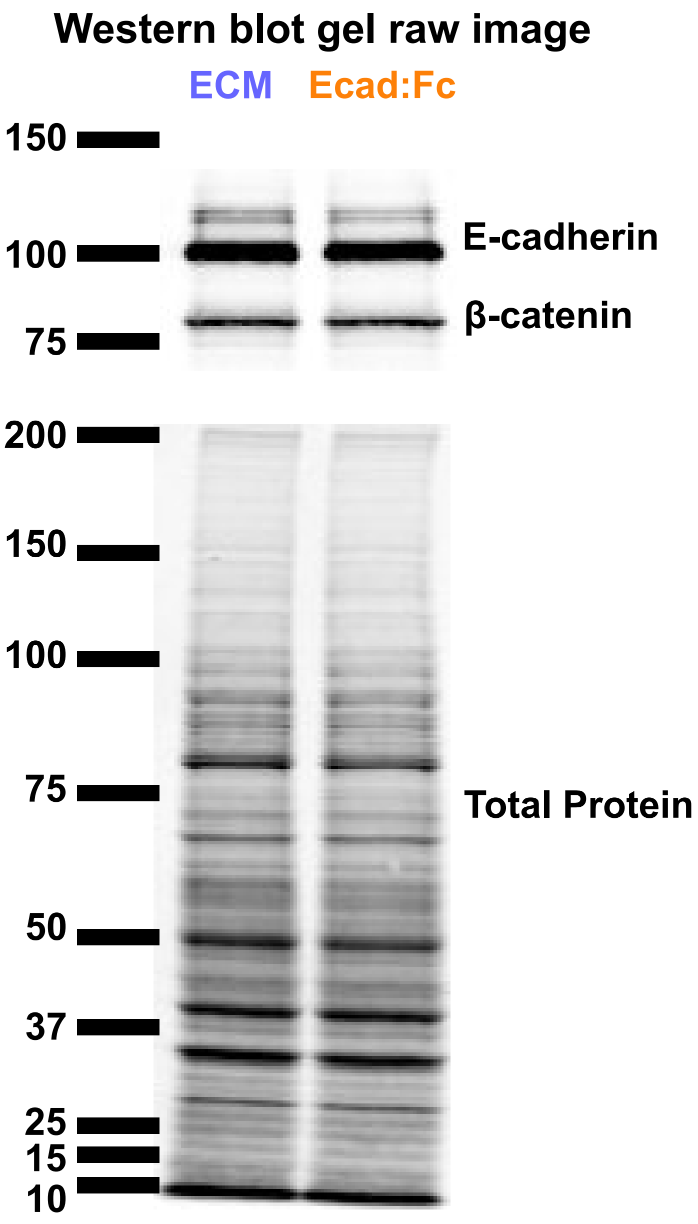


**Fig. S3.** Western blot raw image against E-cadherin and $\beta$-catenin. (Top) To normalize the protein signal with the total protein content, total protein stain was performed. (Bottom)

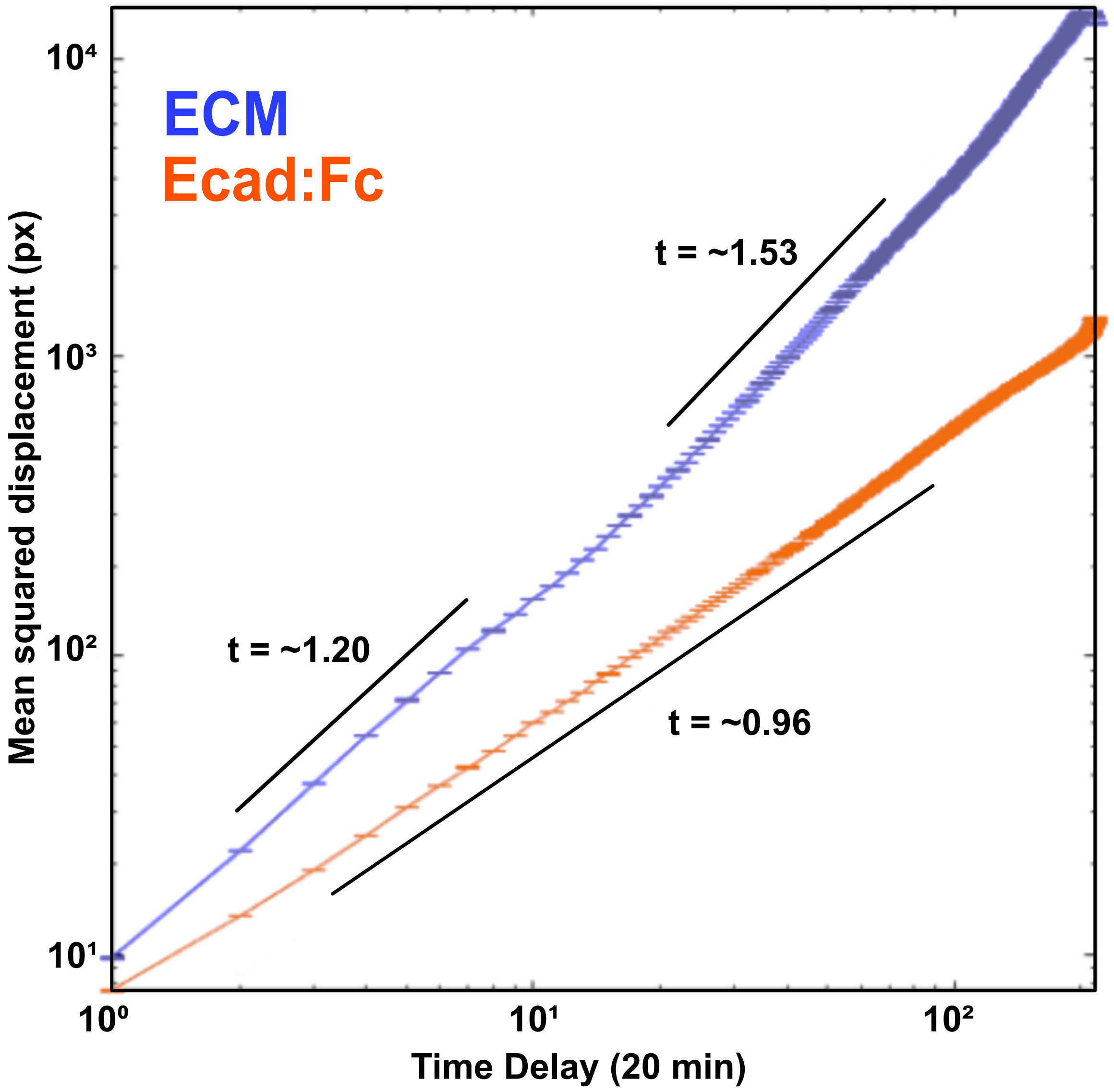


**Fig. S4.** MSD analysis of cell migration on different substrates. MSD of MDCK cells on a collagen substrate shows super-diffusive (or ballistic) motion in two regimes, with increasing diffusivity at longer timescales. On an Ecad:Fc (cell-mimetic) substrate, cells exhibit purely diffusive motion.

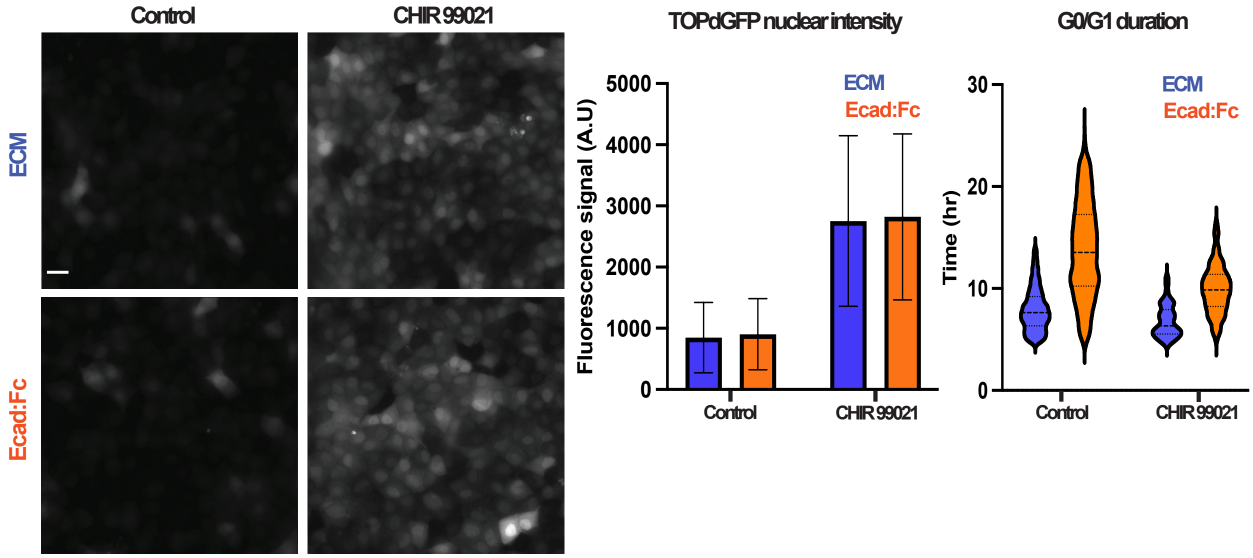


**Fig. S5.** Left: Representative TOPdGFP reporter imaging for WNT activity measurement (Scale bar: 25 $\mu$m). Middle: TOPdGFP nuclear intensity quantification. Right: G0/G1 duration measurement of control tissue and CHIR 99021 treated tissue.

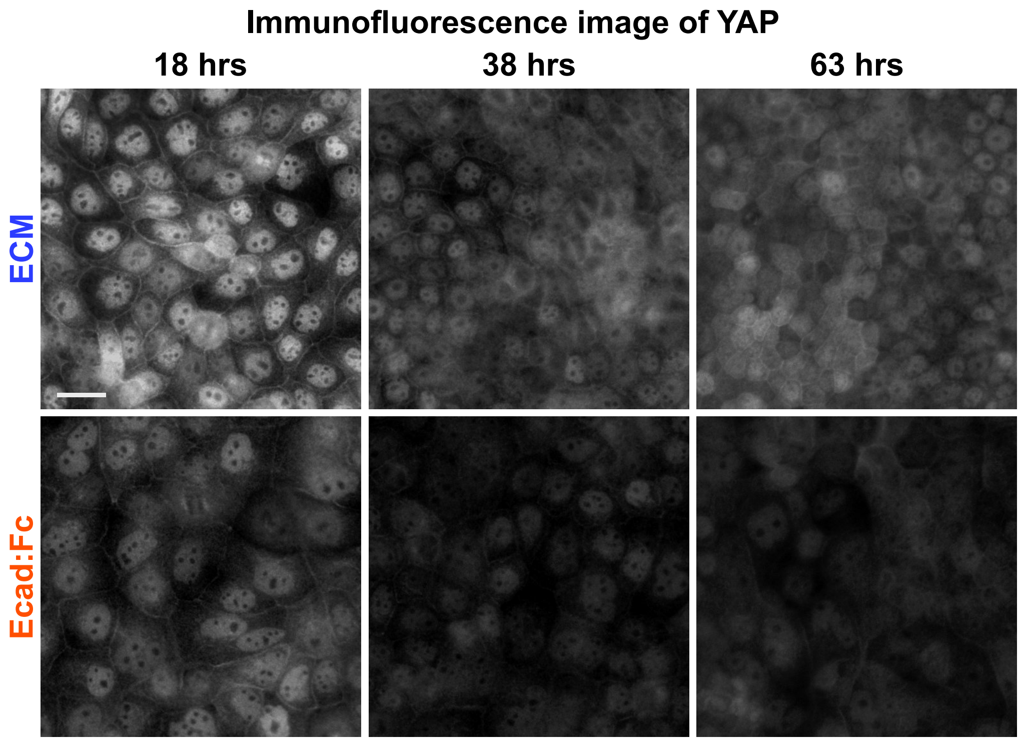


**Fig. S6.** Representative immunofluorescence imaging for YAP at selected timepoints after media flood. (Scale bar: 25 $\mu$m)


Kevin Suh*, Youn Kyoung Cho*, Isaac B. Breinyn, Daniel J. Cohen

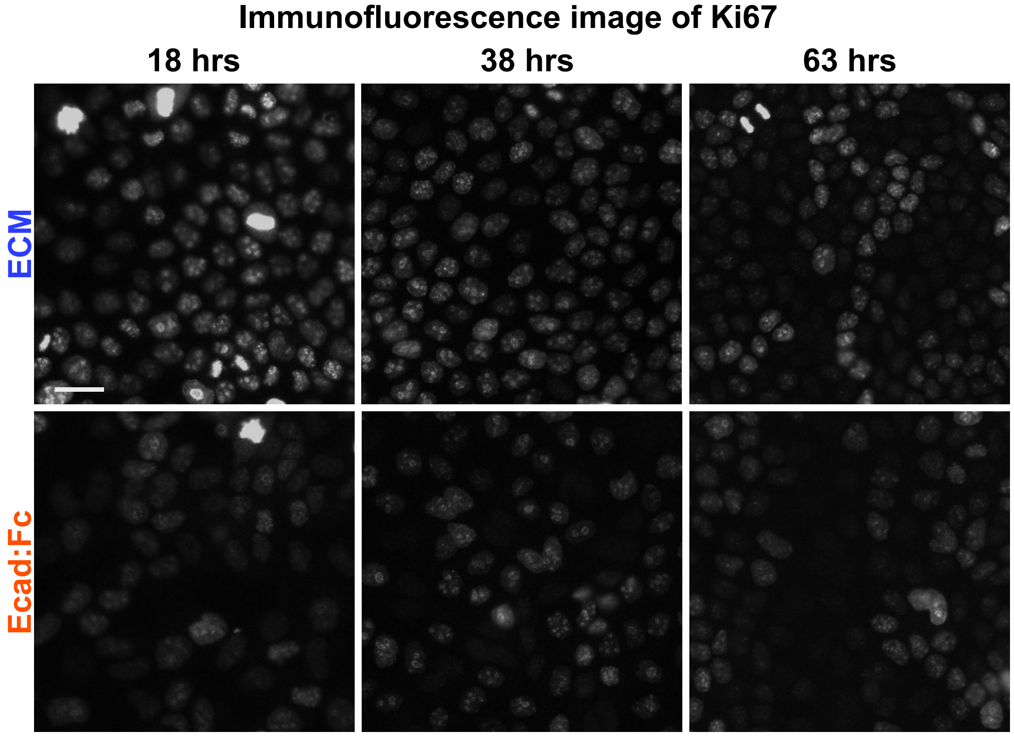


**Fig. S7.** Representative immunofluorescence imaging for Ki67 at selected timepoints after media flood. (Scale bar: 25 $\mu$m)

**Movie S1. Fluorescence E-cadherin imaging of the Ecad:dsRed MDCK monolayer on the Janus substrate (left: ECM side, right: Ecad:Fc side)**

**Movie S2. Phase time-lapse of the MDCK monolayer on the Janus substrate (left: ECM side, right: Ecad:Fc side), time hr:min**

**Movie S3. Phase imaging of the MDCK monolayer on the substrate with thin strip of Ecad:Fc (white lines in the first frame) between ECM on either side**

**Movie S4. Time-lapse of the strain rate calculated from the monolayer on Janus substrate.**

**Movie S5. Fluorescence FUCCI cell signaling marker imaging of the MDCK monolayer on the Janus substrate (left: ECM side, right: Ecad:Fc side; Green: Geminin, Magenta: cdt1), time hr:min**

**Movie S6. Fluorescence FUCCI cell signaling marker imaging of the MDCK monolayer on the substrate with 1 mm strip of Ecad:Fc between ECM on either side, time hr:min**
